# Entorhinal cortex receptive fields are modulated by spatial attention, even without movement

**DOI:** 10.1101/183327

**Authors:** Niklas Wilming, Peter König, Seth König, Elizabeth A. Buffalo

## Abstract

Grid cells have been identified in the entorhinal cortex in a variety of species and allow for the precise decoding of position in space (1–7). Along with potentially playing an important role in navigation, grid cells have recently been hypothesized to make a general contribution to mental operations, including remembering the past and thinking about the future (8,9). A prerequisite for this hypothesis is that grid cell activity does not critically depend on physical movement. Directed attention, which contributes to virtually all mental operations and can be separated from physical movement provides a good test case to investigate this hypothesis. Overt attention in the form of fixational eye movements leads to grid-like firing fields in the monkey entorhinal cortex (3). Here we show that movement of covert attention, without any physical movement, also elicits spatial receptive fields with a triangular tiling of the space. In monkeys trained to maintain central fixation while covertly attending to a stimulus moving in the periphery we identified a significant population (20/141, 14% neurons at a FDR<5%) of entorhinal cells with spatially structured receptive fields. Further, we were able to identify a population of neurons that were labeled as grid cells on an individual basis. This contrast with our recordings obtained in the hippocampus, where grid-like representations were not observed. Our results provide compelling evidence that neurons in macaque entorhinal cortex do not rely on physical movement. Notably, these results support the notion that grid cells may be capable of serving a variety of different cognitive functions and suggest that grid cells are a versatile component of many neural algorithms.

## Introduction

Spatial representations, in the form of place cells and grid cells, have been identified in rodents (1,10), bats (2,11), macaque monkeys (3) and humans (4,5,12). The hippocampal formation, however, also contributes to the processing of memories (13), and conceptual similarities between spatial navigation and the processes involved in remembering and planning suggest that grid cells might support cognitive functions besides spatial navigation (8). This idea resonates well with the recent demonstration that grid cells are active during the exploration of images with eye-movements, i.e. by overt attention (3). These findings revealed that grid cells do not only track an animal's location in space, but can also represent the gaze position of the animal. Shifts of gaze location usually correspond to shifts in attention (14,15), and the neural substrate for overt and covert attention is largely overlapping (16). Accordingly, grid cells in the entorhinal cortex are potentially capable of representing not only gaze location but the locus of attention in general. Here, we investigated whether firing fields of entorhinal cells are activated by movements of covert attention, in the absence of any physical movement. Attention functions as an important control for mental processes (17); accordingly, the hypothesis that grid cells participate in a range of cognitive functions predicts that grid cells may be activated by movements of attention. Here we tested this prediction by recording from single units in the entorhinal cortex in monkeys trained to perform a task of covert attention.

### Results

To examine spatial representations of cells in the entorhinal cortex during the movement of covert attention, we recorded the neuronal activity of 141 neurons in the entorhinal cortex of two rhesus macaque monkeys performing a covert attention tracking task. The monkeys were trained to maintain central fixation while attending to a small (1 °) dot that moved around the computer monitor (see Figure 1). The task of the monkey was to respond by quickly releasing a response bar when the dot changed color. The color change occurred after a random time interval (700-2000ms) following trial onset, and over the course of a recording session, the dot traced out several Hamiltonian cycles, thereby ensuring that in the limit of infinitely many cycles each screen location was visited equally often. Monkeys performed the task at >70% accuracy, with the majority of errors due to saccades to outside of the fixation window. Depending on their motivation, monkeys completed between 2 and 8 cycles per session.

**Figure 1:**
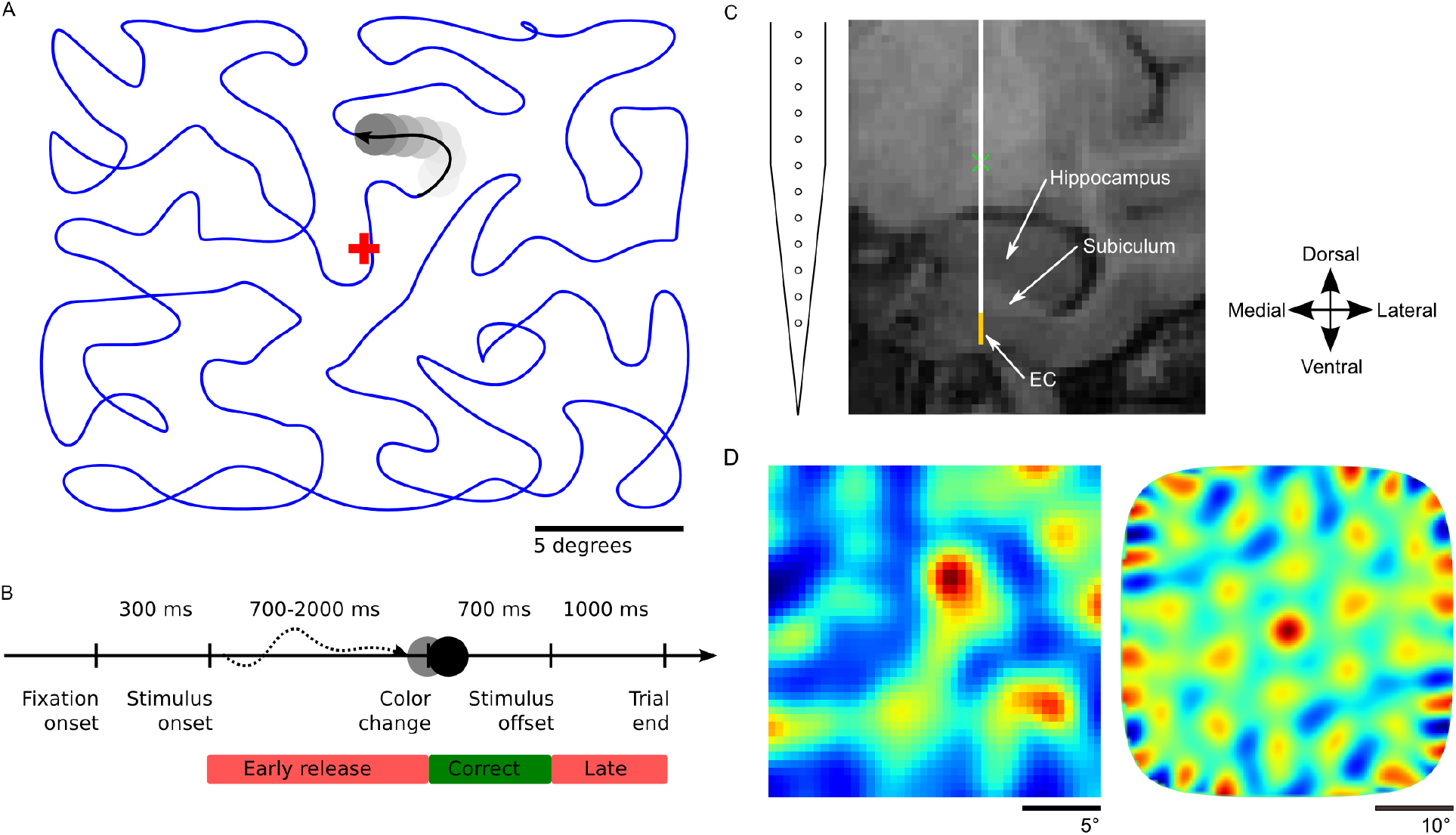
Task design and recording location. **A**, Monkeys were trained to maintain central fixation while a dot moved across the screen and were rewarded for releasing a response bar when they detected a color change of the moving dot. Dot trajectories were smoothed Hamiltonian cycles through a 23×23 grid and covered the screen without a central bias. The blue trajectory was not shown during the experiment. **B**, Temporal structure of a trial. **C**, Example of MRI-guided electrode placement (coronal section through the EC). Recordings were carried out with a 12-site laminar electrode array mounted on a tungsten microelectrode (AXIAL Array, 30μm diameter, 150μm spacing, FHC Inc.). Tungsten electrode is represented in white, the yellow strip represents the span of the recording contacts. EC = Entorhinal Cortex. **D**, Example cell recorded under conditions of covert attention. Left panel shows the estimated firing field (max. firing rate 1.47Hz). The 2D autocorrelation in the right panel shows six peaks surrounding the center which is characteristic for grid cells (gridness score 1.80).

In order to examine spatial representations among entorhinal neurons, we evaluated the standard gridness measure (18–20) along with a novel analysis of the depth of firing rate modulation within the firing fields. Grid cells possess a regular firing field with increased firing rates at the nodes of equilateral triangles that evenly tile space. Importantly, tiling space with equilateral triangles leads to 2D autocorrelation functions with six peaks arranged in a hexagonal structure around the center peak. Whether or not a cell's firing field shows this regular pattern is usually evaluated by quantifying the 60° rotational symmetry of the 2D autocorrelation (18–20). An important property of the gridness score is that the autocorrelation normalizes away the absolute firing rate modulation over space. That is, a cell that shows only a weak change in firing rate as a function of space might have the same gridness as a cell that shows a strong change in firing rate. The gridness score is therefore susceptible to noise, and homogeneous firing fields might produce high gridness scores by chance. Furthermore, for downstream read out of the grid signal the size of firing rate modulation, e.g. the signal to noise ratio is relevant. To better differentiate between noisy and grid-like firing fields, we computed for each cell an index of firing field modulation from its estimated rate map and a gridness score from its 2D autocorrelation function (see Methods).

To assess the statistical significance of spatial representations in our recorded neurons, we computed the same indices on simulated cells with grid, place, or homogeneous firing fields. Cells were simulated by creating spike-trains with an inhomogeneous Poisson (IP) process (Figure 3). The rate function of the IP process was determined by the dot’s trajectory through a firing field and a noise parameter that varied the influence of noise for each simulated cell (see Methods). Simulated cells with a homogeneous firing field therefore control for any potential grid-like structure in the dot's trajectory. The resulting joint distributions of gridness scores and firing field modulation indices represent the likelihood of observing particular patterns of results. To differentiate grid cells, we computed the log likelihood ratio for comparing grid cells with place cells and homogeneous cells (Figure 2 and Methods). The resulting log likelihood ratio expresses how much more likely a specific gridness score and firing field modulation index combination are produced by a grid cell compared to homogeneous or place cells. To control our false-discovery rate we determined the appropriate threshold for classification performance on simulated non-grid cells. Out of 141 recorded neurons, we identified 20 cells (14%) with a log likelihood ratio of > 4.25 deciban (corresponding to a false discovery rate <5%) in favor of cells with spatially structured firing fields according to the two criteria of modulation index and gridness score. Cells classified as grid cells had a lower mean firing rate (0.47Hz) compared to those classified as other cells (1.22Hz), and higher average gridness (1.29 vs. 0.51) and firing field modulation index values (0.1 vs. 0.04). We also tested more conservative thresholds of 10 and 20 deciban (false discovery rates of 1.26% and 0.07%, respectively), which require more evidence to classify a cell as a grid cell. Regardless of the threshold used, we were able to identify a population of neurons that were labeled as grid cells on an individual basis with high significance (X^2^(0, N=141, FDR=0.05)=25.0, p<0.0001, 20, 9 and 3 cells w.r.t. to false discovery rates for different thresholds). We also identified candidate cells that qualitatively resembled grid cells but which could also be explained by assuming a noisy place like firing field (see Figure 2, examples in the lower right).

**Figure 2.**
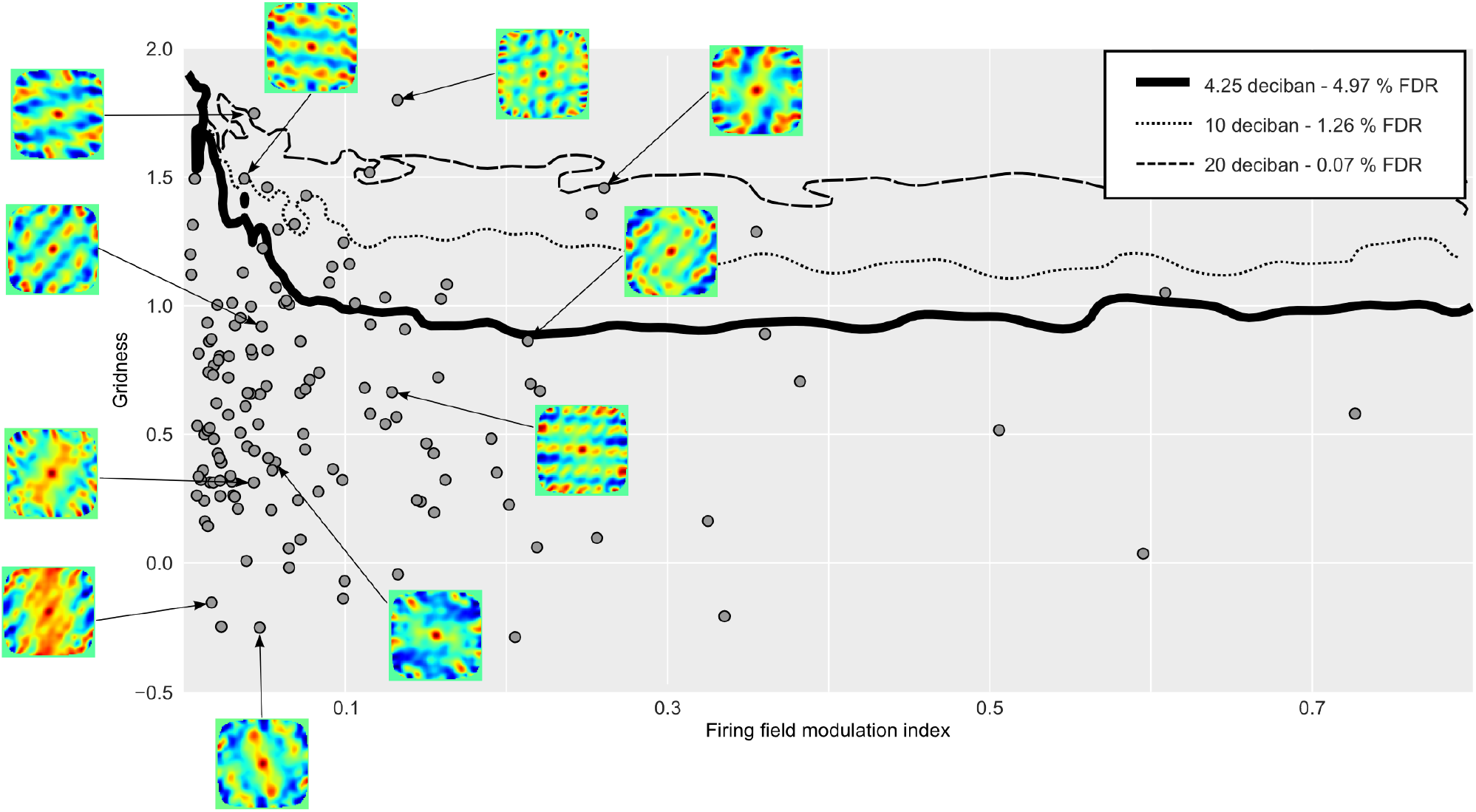
Classification of grid cells. Gray filled circles indicate gridness and firing field modulation index values for our recorded EC units. Solid curves show contours of the log likelihood ratio (in deciban) that indicates evidence in favor or against a cell being a grid cell. 4.25 deciban (thick line) corresponds to a false discovery rate (FDR) of <5% in our simulations, 10 deciban (10:1 odds ratio, thin dotted line) to a FDR of 1.26% and 20 deciban (thin dashed line) to a FDR of 0.07%. The FDR indicates how many simulated non-grid cells are classified as grid cells with each log likelihood ratio value. Cells above the curves are likely grid cells. Insets show the 2D autocorrelation for example cells.

We furthermore sought to validate our novel statistical approach with two different control analyses. First we recorded neuronal activity of 307 neurons from the hippocampus in monkeys performing the same task. There are currently no reports of grid cells in the hippocampus and these neurons can therefore serve as a baseline to determine our empirical false discovery rate. Classifying these cells with the same measures and the 5% FDR threshold yielded an empirical false-discovery rate of 8.5%. The FDR is slightly higher than the FDR derived from simulations because the hippocampal recordings likely contain a different ratio of spatially structured (e.g. place cells) to spatial noise cells. Yet, in the entorhinal cortex (see above) we find significantly more grid-cells than expected from a FDR of 8.5% (X^2^(0, N=141, FDR=0.085) = 5.9, p=0.016; p<0.05 for classifying with the 10 and 20 deciban thresholds as well). Second, we used a bootstrapping analysis to create a joint distribution of indices when the association between firing rate and the dot’s position is destroyed. To this end, we estimated the firing rate of each neuron along the dot’s trajectory and shuffled trajectory segments of 50ms length (700 permutations). We again computed both indices and found an empirical false discovery rate of 5.5%. These control analyses suggest that our main analysis provides an adequate control for the false discovery rate.

We calculated two further controls to eliminate the possibility that our results could be due to the specific set of trajectories used in each session. First, combining spikes generated by different firing fields destroys the spatial structure of the individual firing fields. Any residual grid-like spatial structure of a pooled unit is therefore due to the specific stimulus conditions. We therefore pooled all neurons in each recording session and calculated gridness scores and firing field modulation indices. None of these pooled units was classified as having a grid-like firing field (average and max gridness score 0.36/0.96, average and max firing field modulation 0.02/0.11). Second, we investigated the spatial periodicity of trajectories in isolation. For each recording session, we pooled all trajectories and computed gridness scores and firing field modulation indices by assuming an equal firing rate at all locations. None of these units was classified as a grid cell (average and max gridness score 0.1/1.17, average and max firing field modulation 018/0.033). These controls demonstrate that our results are not due to an inherent grid-like structure in the trajectories.

An important caveat is that our experimental conditions are most sensitive for firing fields with a size that allows multiple repetitions of the grid on the screen. Firing fields that are larger might appear like place or homogeneous cells, and firing fields with spacing smaller than the spacing of our trajectories will appear as spatially structured noise. We can therefore make no definitive statements about the percentage of grid cells in the entorhinal cortex, and our results likely represent a conservative estimate.

## Discussion

In summary, our results provide strong evidence that the activation of cells in entorhinal cortex with spatially structured firing fields does not require physical movement through an environment. Previous results demonstrated that, in stationary monkeys, grid cells can be activated by eye movements as monkeys visually explore complex scenes (3). Here our findings suggest that even eye movements are not required to activate cells with grid like firing fields, and that firing fields in the entorhinal cortex cells can represent the location of attention. That spatial representations in the hippocampal formation can represent locations other than the animal’s current position has been shown in rodents through the identification of theta phase precession (21) and the demonstration that ensembles of hippocampal place cells fire in sequence within a theta cycle, representing places in front of and behind the animal (22–24). More recent studies have shown that sequential firing of hippocampal place cells can represent anticipation of upcoming locations (25), future possible paths when the rat is at a choice point (26,27), and even behavioral trajectories towards a remembered goal in a 2D environment (28). While it is difficult in rodents to assess the location of attention distinct from the location of the animal, these data are consistent with the idea that hippocampal sequences may reflect the animal’s attention being directed along a specific spatial trajectory. Paying attention to the just-completed trajectory could serve to enhance encoding of that experience. Correspondingly, directing attention to future possible paths could serve as a retrieval mechanism to enable optimal choice behavior. These findings also resonate well with a report of grid-like six-fold symmetric BOLD activity while participants navigate a purely abstract concept space (9,29,30) The data presented here demonstrate that entorhinal cells with grid like firing fields can similarly encode information apart from the animal’s current location. Accordingly, these data support the concept that grid cells might implement a structure for mental time travel in the service of episodic memory encoding and retrieval (8).

## Methods

### Summary

Each recording session consisted of the presentation of several cycles of the moving dot. Cycles were smoothed Hamiltonian cycles through a 23x23 grid with 1° spacing and therefore showed no center bias. Each cycle consisted of an average of 114 trials and took the monkeys an average of 13 min to complete. The speed of the dot changed as a function of the trajectory’s curvature, slowing down during corners and speeding up during straight parts. The average speed of a typical trajectory was 1.95 °/s, with a maximum of 3.75 °/s. Each trial ended with the correct detection of the color change, a missed detection, or when the monkey looked outside of the 4.5°x4.5° central fixation window. In subsequent trials, the dot reappeared at its ending location from the previous trial. After the completion of a cycle the next cycle was chosen randomly from a set of 100 precomputed trajectories. Monkeys performed the task at >70% accuracy, with the majority of errors due to saccades to outside of the fixation window. We only analyzed data from trials with correct detection of the color change and constant fixation within the fixation window. Depending on their motivation, monkeys completed between 2 and 8 cycles per session. The monkeys were head-fixed and seated in a chair such that the monitor's center corresponded to their neutral eye position. Spikes were recorded (40kHz) from the entorhinal cortex with a laminar electrode array mounted on a tungsten microelectrode (12-site, 150-μmm spacing; FHC). Rate maps were computed with a standard Gaussian smoothing procedure(3,20).

Gridness scores were calculated using standard equations (3,18–20), but we additionally validated that our implementation was highly accurate in distinguishing grid from non-grid cells in simulated data (see “Gridness score implementation”, “Unit simulation”, “Log likelihood ratios and classification of units” and methods Figure 3, 4 and 5). We computed a firing field modulation index to separate spatially structured firing fields from homogeneous noise-dominated fields. The firing field modulation index captures how much of the spatial variation in firing rate can be explained by the mean firing rate over space:

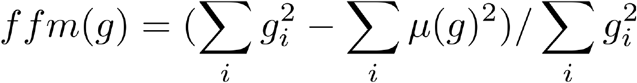

where g is the 2 dimensional rate map which is linearly indexed by i and μ(g) is the mean firing rate. Significance was established by computing the logι_0_ likelihood ratio between the likelihood that a specific cell’s gridscore and firing field modulation index was produced by a noisy grid-cell or a cell with a noisy homogeneous or noisy place field like firing field (see “Log likelihood ratios and classification of units”). All experiments were carried out in accordance with protocols approved by the Emory University and University of Washington Institutional Animal Care and Use Committees.

**Figure 3.**
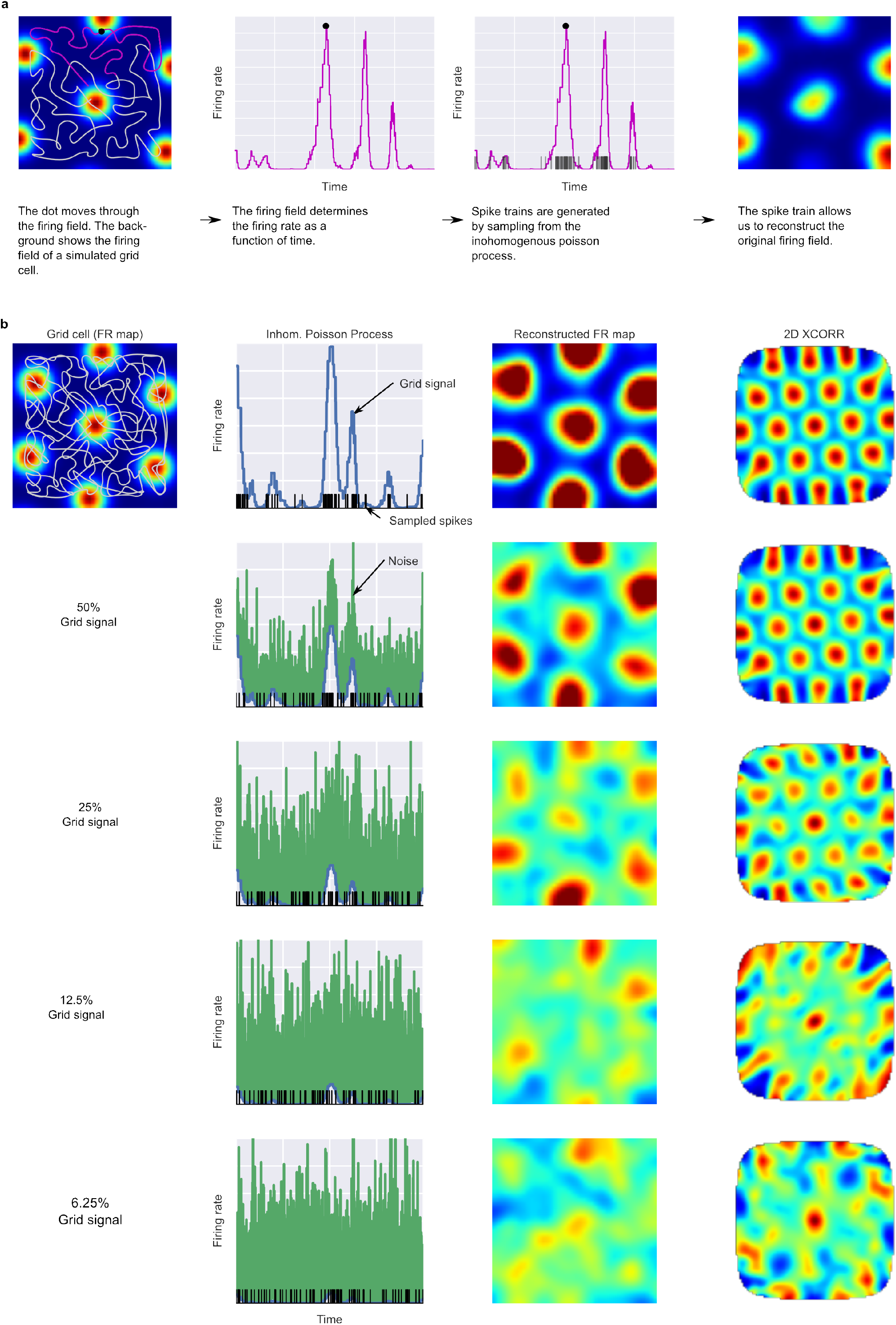
Simulation of units from grid-like firing fields. Panel **A** shows how the dot’s trajectory and a 2D firing field generate the time varying firing rate for a simulated unit. Panel **B** shows how different amounts of noise influence the reconstructed firing fields and 2D autocorrelations.

### Behavioral task

We recorded neuronal activity in the entorhinal cortex in one male and one female monkey (maccaca mulatta) trained to perform a covert attention tracking task. The task required the monkeys to attend to a dot as it moved across the screen. The monkey's task was to maintain central fixation within a 4.5 ×4.5 degree window and to release a bar when the moving dot changed color. Gaze location was monitored with an infrared eye-tracking system (ISCAN, Inc.). Each trial began with the onset of a central fixation cross. After the monkey fixated the cross for 300ms, a moving dot appeared. The dot moved for a random duration between 700 and 2000ms before it changed color (from light gray to white). After the color change, a bar release that occurred within 700ms was counted as a successful detection. Early and late bar releases as well as fixational breaks were counted as failures. Upon bar release or 700ms after the color change the moving dot disappeared. The monkey was rewarded with a food-based slurry on successful trials. In the next trial, the dot continued on the same trajectory, e.g. started where it had disappeared in the previous trial. Monkeys worked for as long as they were well-motivated and completed between two and eight cycles per session.

The dot followed trajectories that visited different parts of the screen equally often. Dot trajectories were Hamiltonian cycles through a grid of 23x23 nodes (with 1° spacing) that evenly covered an area of -11:11° of visual angle on the screen. Each trajectory was created by using the Concorde (http://www.math.uwaterloo.ca/tsp/concorde/) traveling salesman problem solver to find a path that visits each node of the 23×23 grid exactly once. The x and y coordinates of the trajectory were smoothed with a Gaussian kernel with σ=60° to yield a smooth trajectory. The smoothing also ensured that the dot continuously changed its speed (mean for a typical trajectory 1.95°/s, 3.75°/s maximum). At the beginning of a recording session and after completion of each cycle the next trajectory was chosen randomly from a precomputed set of 100 trajectories. Stimuli were displayed on a 19” CRT monitor with a refresh rate of 120Hz. The monkeys were head-fixed with a distance of 60cm between eyes and monitor.

### Electrophysiological recordings

Spikes (40Khz) were recorded from the EC with a laminar electrode array mounted on a tungsten micro electrode (AXIAL array: 12-site, 30μm diameter, 150μm spacing, FHC, Inc.) with recording hardware from Plexon Inc. Subsequent analysis was carried out with custom MATLAB (The MathWorks, Natick, MA) and python code. Recordings from monkey MP (male) were carried out in the right hemisphere and recordings from monkey PW (female) were in the left hemisphere. Recording sites were planned with the help of MRI scans of each monkey and an atlas. Recording sites were located in the posterior part of the entorhinal cortex and close to the rhinal sulcus. Each recording started by slowly lowering a stainless steel guide tube containing the laminar array through a craniotomy. The array was then slowly advanced to its final position while the monkey conducted unrelated tasks. Recordings were started a few minutes (~10) after the array reached its final position to allow the tissue around the electrode to settle. Spikes were sorted using offline methods (OfflineSorter, Plexon Inc.). We discarded units that did not show stable firing behavior through an entire cycle. In total we analyzed 69 units from monkey MP and 72 units from monkey PW

Neural recordings from the hippocampus were conducted in a similar manner. Four independently moveable tungsten microelectrodes (1 to 3 MΩ, FHC Inc.) were lowered down into the hippocampus with the use of coordinates derived from MRI scans. Recordings took place in the left anterior portion of the hippocampus in one monkey (PW) and in the right posterior portion of the hippocampus in another monkey (TO). Neurons were recorded from CA1-CA4 subfields, dentate gyrus, and subiculum. For the hippocampal recordings, we recorded from the covert attention task for 2 to 6 cycles per session following a ~1 hour recording during which monkeys freely viewed images.

### Rate maps and autocorrelation

Rate maps and autocorrelation functions were computed analogously to1. We computed 2D histograms of spike and dot positions with bins of size 0.5°×0.5°. To estimate a firing rate map we smoothed both histograms with a Gaussian kernel (σ=1.25°) and divided the spike counts by the amount of time the dot spent in the respective locations. 2D autocorrelations were computed by shifting the firing rate map relative to itself and computing the coefficient of correlation for the overlapping maps.

### Statistical analysis

Our statistical analysis proceeded in several steps. Our goal was to classify the reconstructed firing fields of recorded units as grid like or not grid like. In a first step, we identified two measures of firing behavior that were suitable for this classification. In a second step, we estimated the joint distribution of these measures when the generating cell is either a noisy grid cell, a cell with a noisy homogeneous firing field or a noisy place field like cell. This allowed us to compute a log likelihood ratio that indicates whether or not a firing field is caused by a grid cell or by the other types of simulated cells. We then used the log likelihood ratio for classification of cells as grid cells or other cells.

Both of these steps made extensive use of spike trains of simulated neurons. We therefore first describe our model for simulating spike trains and then describe the classification process.

### Unit simulation

We simulated units by assuming that firing behavior is the result of an inhomogeneous Poisson process (IP). This process allows to model different firing rates for different positions of the moving dot. The rate function of the IP is determined by a firing field, through which the dot passes, and a noise component. Sampling from the IP defined by this rate function yields artificial spike trains caused by the underlying noisy grid field. We simulated spike trains from noisy grid-like, noisy homogeneous or noisy place field-like firing fields. Grid-like firing fields were created by placing 2D Gaussian distributions at the nodes of a grid that tiles space with equilateral triangles:

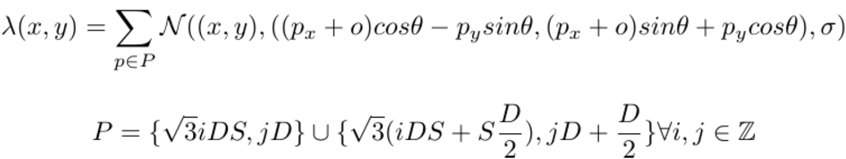

where λ(x,y) is the firing rate of the grid field at location (x,y), D is the distance between peaks, S is a scaling parameter, θ rotates the grid field, o shifts the grid field and σ specifies the size of each grid node. The grid component of the rate function represents the firing rate in the firing field along several randomly drawn dot trajectories. The dot trajectories transform the 2D firing field into a 1D time-and space-varying firing rate. Simulated units were evaluated with the same pool of trajectories used for the recorded units. The noise component of the rate function is drawn from an exponential distribution (rate=1) for each location in the trajectory. The amount of noise and grid signal in the time and space varying firing rate is governed by the parameter β_signal_:

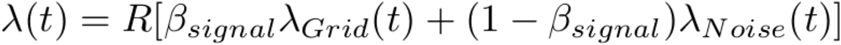

λ_Grid_, λ_Noise_ are the grid and noise dependent components of the firing rate and are scaled to have an expected firing rate of 1. 0 < βsignal < 1 specifies how much the grid signal contributes to the firing rate. R is the desired mean firing rate (here uniformly sampled from the empirical firing rates). λ(t) is therefore the firing rate of the IP. Spike trains were then generated by producing a sample from the IP defined by the time-and space-varying firing rate λ(t). The rate function has a temporal resolution of 120Hz. Methods Figure 3 shows a graphical overview of unit simulations.

We simulated 5000 grid cell units for each of 9 different β_signal_ values (i^2^ for evenly spaced i between 0.18 and 1). All other parameters were drawn from random distributions:

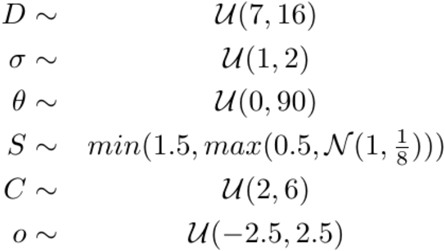

Where C is the number of dot trajectories used for the simulation. Parameters are given in degrees of visual angle and ranges were chosen to allow broad coverage of potential grid-like firing fields. We discarded all firing fields that had less than 3 peaks in the area covered by the trajectory; these are a priori not recognizable as grid-like. This resulted in 27,450 units whose firing rate was determined by a grid-like firing field. We also simulated units based on spatially structured firing fields that were non grid-like. We used place-like firing fields that had only one peak by setting the spacing parameter of the grid-like firing fields to larger than twice the screen size. The standard deviation was randomly drawn from the interval [3,4] to allow large parts of the trajectory to be influenced by the place field. The scaling was set to 1. We used 15 β_signal_ values (i^2^ evenly spaced i between 0 and 1) and simulated 2500 units with each β_signal_ value. This resulted in 37,500 simulated units whose firing fields ranged from completely uniform (β_signal_ = 0) to spatially structured fields (βsignal > 0). To summarize, we simulated a large range of plausible grid fields that can be resolved by our trajectories. In addition we simulated cells with noisy homogeneous firing fields and cells with a noisy place-like firing field.

### Gridness score implementation

Computation of the gridness score is analogous to (19,20). In this computation, an ellipse is fit around six peaks in the 2D autocorrelation function and the rotational correlation is computed at 30, 60, 90, 120 and 150 degrees. The gridness is then defined as:

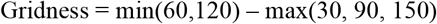

Brandon et al. (20) noted that this measure is sensitive to how the ellipse is fit in the 2D autocorrelation. Choosing the closest six peaks, for example, might include a peak that is due to noise in the firing field and omits a 'correct' peak that is further away. We used a two-step procedure to fit an ellipse. In a first step, potential ellipses that likely contain six peaks and six valleys were identified. This was achieved by predicting for a large space of possible grid fields where the most central peaks and valleys would be in the 2D autocorrelation for this grid field. Grid fields were parameterized by grid spacing, aspect ratio, size of the peaks, rotation of the grid field and shift of the peaks along the ellipse. The prediction r(x,y) was computed according to the following equations:

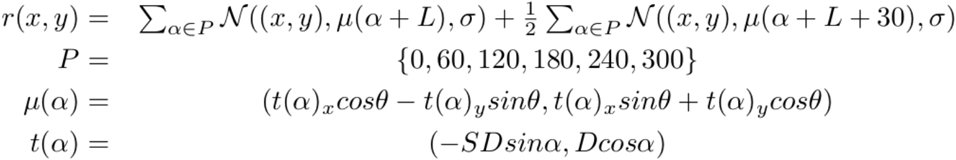

where D is the grid spacing, θ is the rotation of the grid field, σ describes the size of the grid peaks, S relates to the aspect ratio of the grid field and L describes the shift of the peaks along the ellipse. We constrained the set of candidate ellipses to those that passed through at least one non-central peak in the 2D autocorrelation. Methods Figure 4 shows several example predictions. The second step consisted of selecting ellipses that capture structure in the 2D autocorrelation. To this end we computed the RMSE (root-mean-square error) between all ellipse predictions and the 2D autocorrelation functions. We then used a minimum shift algorithm to select ellipses that captured six peaks and valleys in the autocorrelation function better than ellipses in the local neighborhood of possible ellipses (the window of the minimum shift algorithm is N/5). The final gridness score was then the maximum of the gridness scores computed for these ellipses.

**Figure 4.**
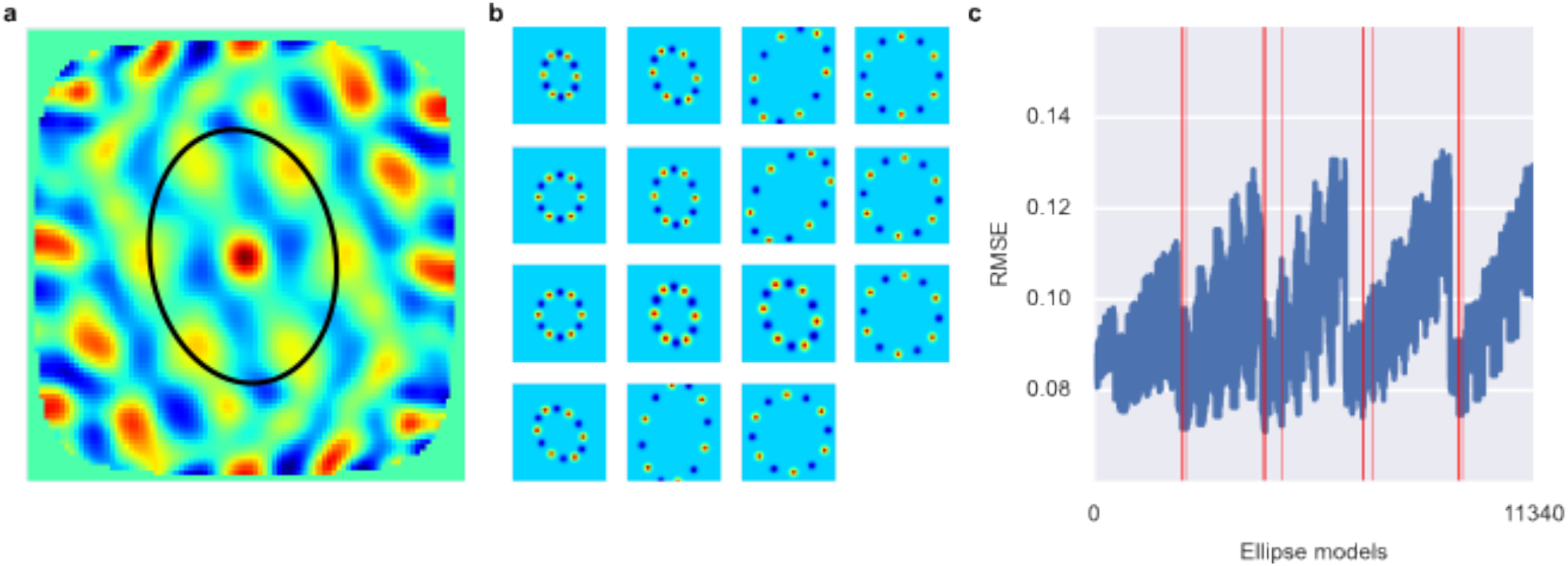
Ellipse fitting on 2D autocorrelation maps. **A** shows one example 2D autocorrelation function and fitted ellipse that passes through six peaks. **B** shows a set of candidate ellipses with hexagonal peaks and valleys. To identify best-fitting ellipses we generated a large number of potential ellipses that passed through at least one peak in the 2D autocorrelation and the computed the RMSE between ellipses and 2D autocorrelation. **C** shows the RMSE of a large number of candidate ellipses, arranged by a linear index into the 5-dimensional parameter space for the ellipses. Red lines indicate ellipses selected by a peak shift algorithm whose maximum gridness score was assigned to this unit.

To evaluate the power of the gridscore implementation to separate grid cells from non-grid cells, we computed gridness scores for all simulated units with grid-like firing fields. We then compared how well the gridscore separated units with firing rates determined by noise (β_signal_ = 0) from those whose firing rate was at least partly determined by the grid firing field (β_signal_ >0). To this end, we computed ROC curves and used the area under the curve as a measure of discriminatory power. Methods Figure 5 shows these ROC curves and the AUC as a function of β_signal_.

**Figure 5.**
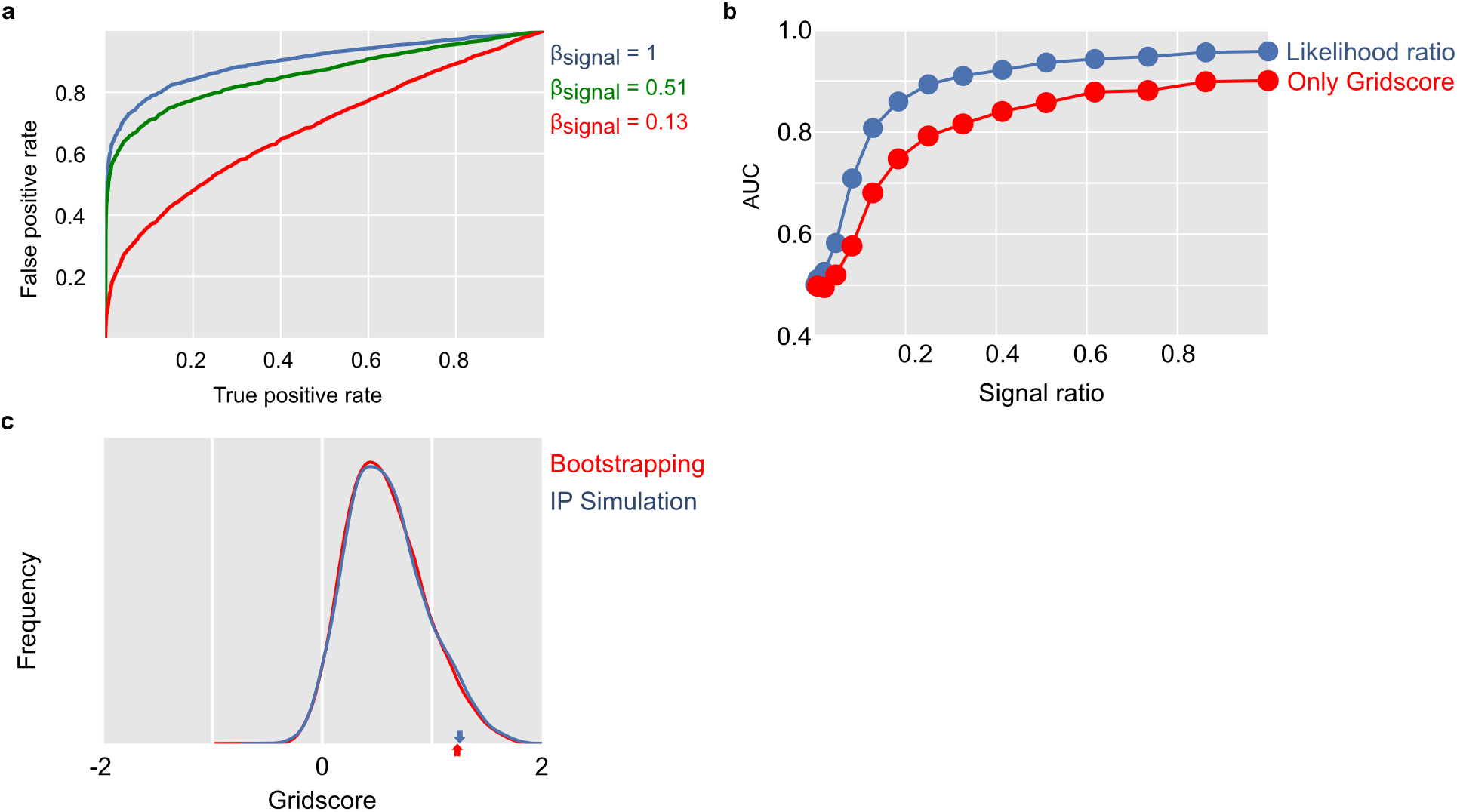
Evaluation of gridness score implementation and log likelihood ratio classification. Panel **A** shows ROC curves for separating simulated grid-like units and simulated units with homogeneous units. Panel **B** compares using only the gridness score with our approach of computing log likelihood ratios based on the ffm and gridness score. Panel **C** compares gridness scores obtained by shuffling spike sequences along the dots trajectory with gridness scores obtained from simulating units with homogenous noise.

### Log likelihood ratios and classification of units

To differentiate grid cells from non-grid cells we computed the log likelihood ratio that compares the evidence that a gridscore and firing field modulation (ffm) combination is generated by a grid cell to the evidence that it is generated by a cell with a homogeneous or place field-like firing field:

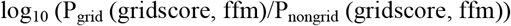

The probabilities Pgrid and Pnongrid were computed from the simulation results with grid-like, place field-like and homogeneous firing fields. We estimated each probability distribution with a kernel density estimator that uses a Gaussian kernel and 10-fold cross validation for bandwidth selection. Since the gridness score and firing field modulation measures are bounded, we applied a logit transformation to avoid boundary artifacts. We only considered units with β_signal_ > 0.15 as simulations of grid cells for the grid density estimate. Smaller β_signal_ values produced firing fields qualitatively close to those obtained with β_signal_ = 0 and thereby inflated the likelihood of observing small gridness scores and firing field modulation values from the grid cell simulation. Removing these units for the density estimation therefore makes our estimates more conservative. The log likelihood ratio expresses how much evidence we have that a particular gridness score and firing field modulation combination is produced by a grid cell relative to the evidence for place cells or cells with a homogeneous firing field.

To evaluate the power of the log likelihood approach to separate grid cells from non-grid cells, we compared how well the log likelihood value separated units with firing rates determined by noise (βsignal = 0) from those whose firing rate was at least partly determined by the grid firing field (β_signal_>0). In analogy to our evaluation of different gridness scores we used the area under ROC the curve as a measure of discriminatory power. Methods Figure 5B shows AUC as a function of β_signal_ in comparison to only using the gridness score. The log likelihood approach performed better than only using the gridness score for all noise levels. Figure 6 shows 2D autocorrelation functions and ratemaps for all EC cells.

**Figure 6.**
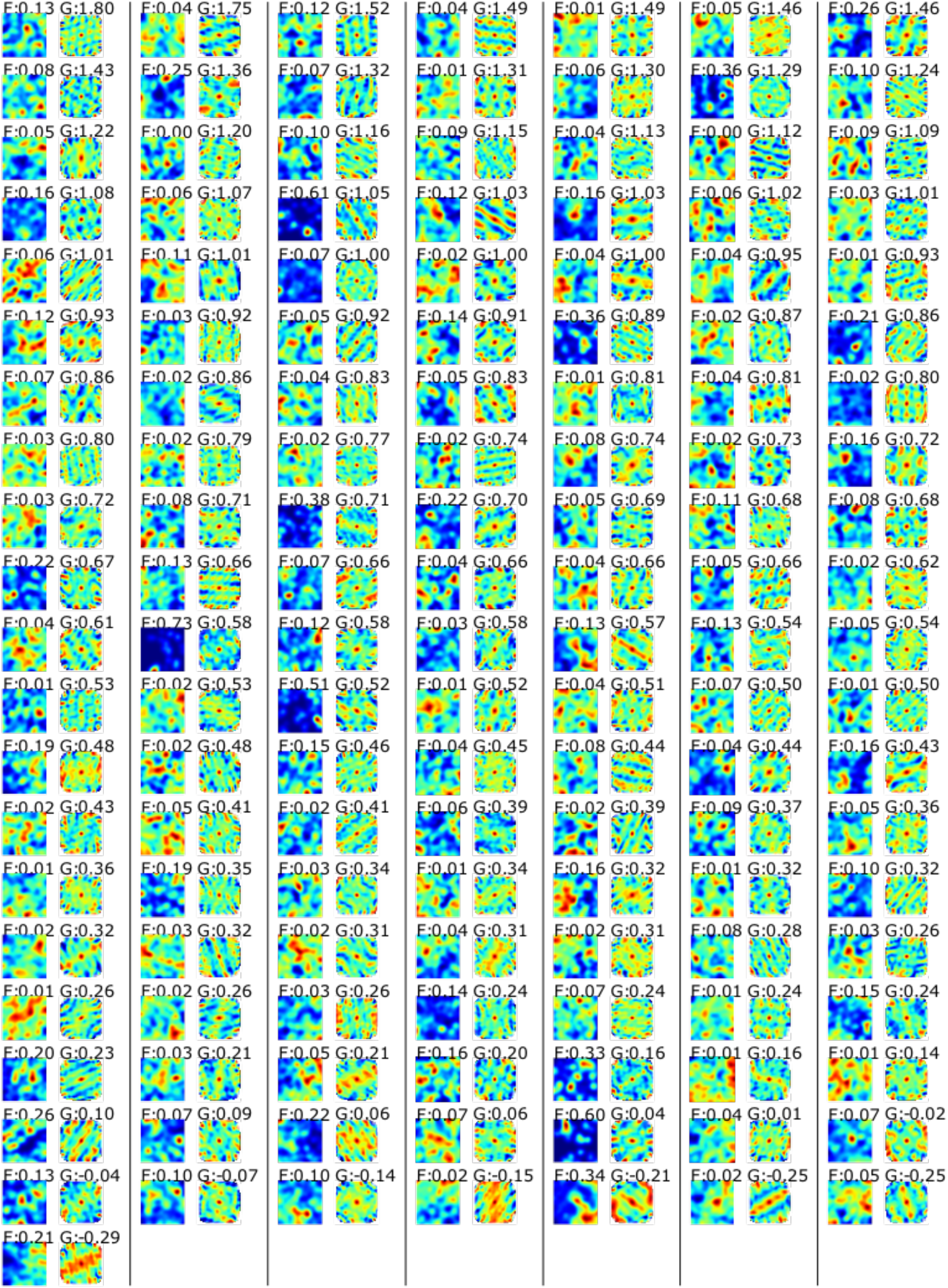
Rate maps (left) and autocorrelation functions (right) for all recorded EC cells. Numbers on top give the firing field modulation index (F) and the gridness score (G).

## Acknowledgements

This study was supported by National Institutes of Health Grants MH080007 (E.A.B.) and MH093807 (E.A.B.), and the Office of Research Infrastructure Programs (ORIP) Grants P51OD010425 (Washington National Primate Research Center) and P51OD011132 (Yerkes National Primate Research Center), the FP7 project eSMCs IST-270212 (N.W., P.K.) and SFB 936 (B6) Multi-Site Communication in the Brain (P.K.).

## Author Contributions

NW collected EC data and analyzed all data. SK collected hippocampus data. All authors designed the study and analysis and wrote the paper.

## Author Information

The authors declare no competing interests.Correspondence and requests should be addressed to NW.

